# Hayai-Annotation v3.0: A functional gene prediction tool that integrates orthologs and gene ontology for network analysis

**DOI:** 10.1101/2024.06.05.597500

**Authors:** Andrea Ghelfi, Sachiko Isobe

**Author notes:** Corresponding Author Andrea Ghelfi, National Institute of Genetics, Yata, 1111, Mishima, Shizuoka, 411-8540, Japan.

## Abstract

Hayai-Annotation v3, an R-package integrated with the R-Shiny browser interface, utilizes two methods for functional annotation: DIAMOND for sequence alignment using UniProtKB Plants as the database, and OrthoLoger, the official OrthoDB tool for ortholog inferences. The GO enrichment accuracy was assessed by a CAFA-evaluator, demonstrating that Hayai-Annotation v3’s accuracy was comparable to that of the benchmark, BLAST2GO. We here propose a method to explore genome evolution and adaptation from a different perspective, by creating networks and heatmaps correlating orthologs with gene ontology (molecular function and biological process) from their co-occurrence tables. This approach enhances the ability to infer functions of uncharacterized genes by associating orthologs with gene ontology terms and the ability to visualize the distribution of gene numbers correlated with co-occurrence patterns across different species. To our knowledge, this is the first attempt to correlate orthologs with GO (MF and BP) to construct a gene network, providing a comprehensive, cross-species view of gene distribution and function. Hayai-Annotation v3 not only retains the convenience of previous versions but also enhances ortholog analysis functionality, allowing for evolutionary insights from gene sequences. Hayai-Annotation v3 is expected to contribute significantly to the future development of plant genome analysis.

## Introduction

The advent of Next Generation Sequencing (NGS) and advances in bioinformatics for genome sequence analysis have led to exponential growth in plant genome information^1^. This surge in genomic data is crucial for accelerating progress in plant research, as it enables molecular biologists to obtain extensive and accurate knowledge of gene profiles in relevant genomes. Numerous tools are available for the annotation of gene and protein functions, such as OmicsBox/Blast2GO^2^, Mercator4^3^, and FunctionAnnotator^4^. These tools employ various algorithms for sequence alignment and Gene Ontology (GO) annotations; for instance, Omics/Blast2GO utilizes BLAST^5^, DIAMOND^6^, and InterProScan^7^. Mercator4 applies BLAST and HMMER3^8^, and FunctionAnnotator employs LAST^9^ and RPS-BLAST^10^. However, these tools often require extensive runtime and do not consider the Protein Existence (PE) level, which represents the evidence validating a protein’s presence.

Meanwhile, with the increasing availability of diverse plant genome sequences, cross-species gene conservation analysis has become more prevalent. Orthology inference is one of the most crucial methods in comparative evolutionary studies, providing accurate predictions of gene functions. Various methods of ortholog detection include OrthoLoger^11^, OrthoFinder^12^, and OrthoMCL^13^. OrthoDB, a leading resource for precomputed gene orthology, implements OrthoLoger. While OrthoDB aspires to encompass all species in order to facilitate accurate comparative research, the current data volume in OrthoDB is approaching the limits of computational resources, as Kuznetsov et al. (2023)^11^ noted. Moreover, not all genes readily fit into ortholog groups using OrthoLoger, prompting reliance on sequence similarity analysis as a standard method for inferring protein functions. This method is particularly prevalent with resources like UniProt Knowledgebase (UniProtKB) as a database, which provides additional annotation layers such as GO, InterPro, PFAM, and PE.

To address these issues, we developed Hayai-Annotation Plants^14^, an ultra-fast, accurate, and comprehensive functional gene annotation system specifically for plants. One of its advantages is the ability to run locally, eliminating the need for users to upload their data to public websites. Hayai-Annotation Plants integrates USEARCH, an algorithm significantly faster than BLAST, for sequence similarity searches against the UniProtKB taxonomy Embryophyta (land plants). It offers five layers of annotation: protein name, GO terms (Biological Process, Molecular Function, and Cellular Component), enzyme commission number, protein existence level, and evidence type.

In this manuscript, we report an updated version, Hayai-Annotation v3, which easily identifies orthologs of plant species. This version implements OrthoLoger to map orthologs to OrthoDB v11 and uses DIAMOND to align sequences against UniProtKB-Plant. It also generates networks and heatmaps correlating orthologs with gene ontology (molecular function and biological process). These approaches aim to enhance the ability to infer functions of uncharacterized genes by associating ODB IDs with GO molecular function and biological process terms, as well as to visualize the distribution of gene numbers correlated with co-occurrence patterns across different species. To our knowledge, this is the first attempt to correlate orthologs with GO (MF and BP) to construct a gene network, providing a comprehensive view of gene distribution and function across species.

## Materials and methods

### Database preparation

The OrthoDB v11 dataset was downloaded from “https://data.orthodb.org/download/odb11v0_all_og_fasta.tab.gz“. The UniProt-Plants dataset, consisting of uniprot_sprot_plants.dat.gz and uniprot_trembl_plants.dat.gz, was downloaded from “https://ftp.uniprot.org/pub/databases/uniprot/current_release/knowledgebase/taxonomic_divisions/“. Taxonomic information was downloaded from ftp://ftp.ncbi.nih.gov/pub/taxonomy/taxdump.tar.gz to identify plant species within the OrthoDB v11 dataset. All downloaded files underwent MD5sum verification.

Orthologs were enriched by extracting sequences of plants species from OrthoDB to create OrthoDB-plants, which served as the primary reference to match orthologs with UniProt-Plants sequences through the utilization of USEARCH^15^ (64-bit), employing a minimum sequence identity of 50%, an E-value of 1e-6, and ensuring 75% query and target coverage (Figure 1A). The information from UniProtKB, such as sequences in FASTA format, accessions, GO, PFAM, InterPro and PE IDs, was extracted from UniProt “dat” files using BioPython scripts. The titles of the sequences from UniProtKB were updated to incorporate GO, PFAM, InterPro, PE, and the newly assigned OrthoDB OGs.

**Figure 1.**
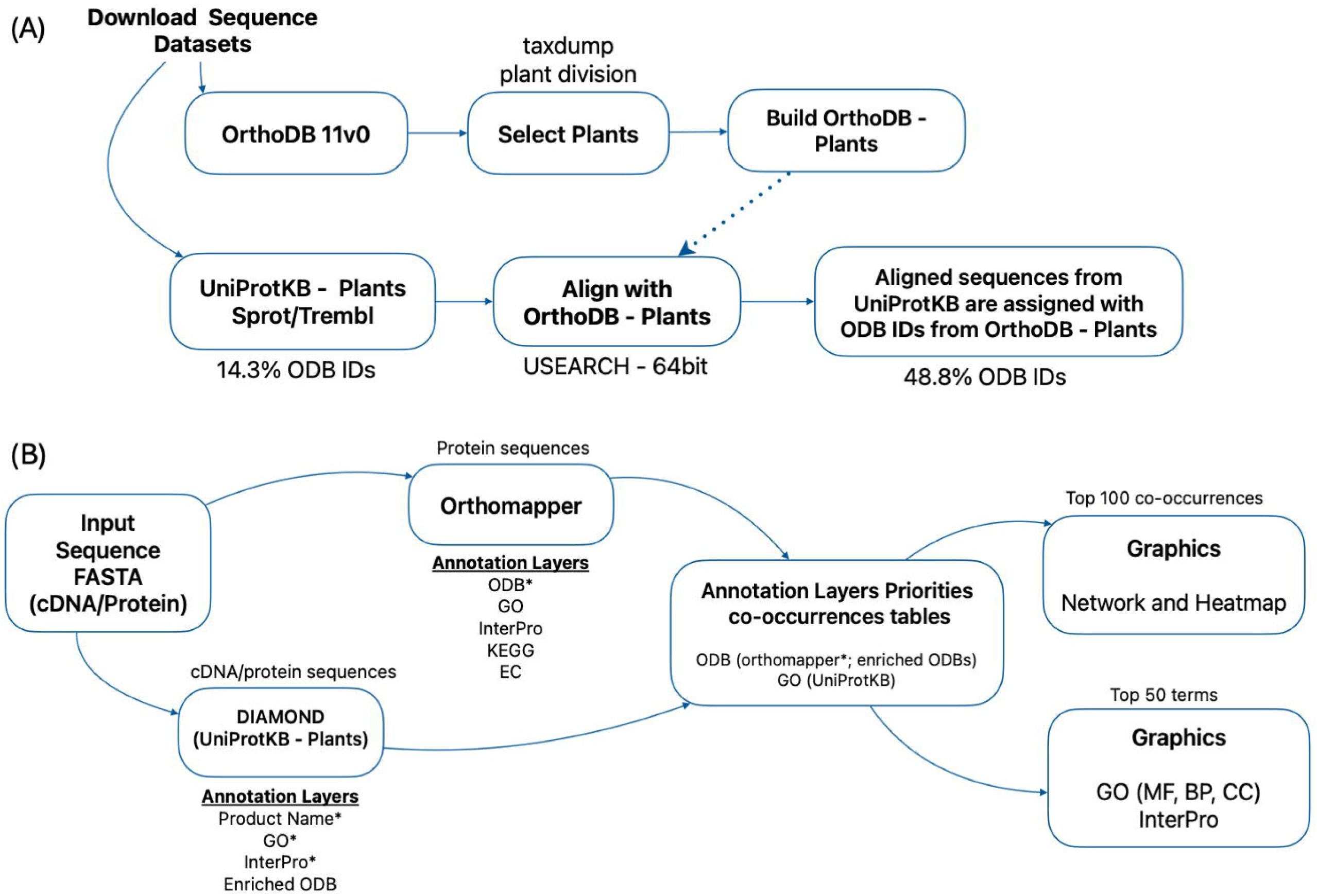
Schematic representation of the pipelines for the (A) ODB-enrichment of UniProtKB by using OrthoDB-plants; (B) tools implemented in Hayai-Annotation (Orthomapper and DIAMOND against UniProtKB-Plants) and in the generation of co-occurrence tables and graphics (networks, heatmaps, and distributions of InterPro and each GO domain).

### Annotation engines

Hayai-Annotation v3 is an R-package employing the R-Shiny browser interface. It uses two methods for functional annotation: DIAMOND (v2.1.9) for sequence alignment using UniProtKB Plants as a database and the orthomapper function from OrthoLoger (v3.0.5, “https://orthologer.ezlab.org/“) using the node Viridiplantae to detect orthologs. Since Hayai-Annotation v3 uses two methods to infer protein function, whenever the query sequence has a hit in both methods, the ODB IDs from orthomapper receive higher priority. They are therefore selected for the subsequent network analysis and the assessment of the co-occurrence of ODB-IDs with GO IDs (MF and BP, analyzed independently) (Figure 1B).

The network and heatmap graphics were generated based on the top 100 co-occurrences of ODB IDs with GO IDs (MF and BP), determined by the number of associated genes. The network graph was constructed using the R packages igraph and ggraph, while the heatmap was created with reshape2 and ggplot2. For the co-occurrence calculations, the source of GO IDs utilized annotation from UniProt-Plants.

### Output files

Hayai-Annotation v3 generates 7 tables (TSV format) and 8 graphics (PDF format, see Supplementary_File_S1 for descriptions). The main table, Hayai_annotation_v3.0.tsv, contains the full annotations from both DIAMOND and orthomapper. Four tables aggregate the results for each GO domain (MF, BP, CC) and InterPro annotation layers. Two tables show the co-occurrences of ODB ID and GO (MF and BP, independently).

The network and heatmap graphics consider the top 100 co-occurrences of ODB ID and GO (MF and BP, independently). The graphics for the distribution of GO (MF, BP and CC) and InterPro consider the top 50 counts for each annotation layer.

### Datasets

The performance of Hayai-Annotation v3 was evaluated using the peptide sequences of

*Arabidopsis thaliana*, domesticated rice, and wild rice as described below.

*A. thaliana:*

https://www.arabidopsis.org/download/file?path=Proteins/Araport11_protein_lists/Araport11_pep_20220914_representative_gene_model.gz

Domesticated rice, *Oryza sativa* Japonica Group, Os-Nipponbare-Reference-IRGSP-1.0: https://rapdb.dna.affrc.go.jp/download/archive/irgsp1/IRGSP-1.0_protein_2024-01-11.fasta.gz Wild-type rice, *Oryza punctata*:

https://ftp.ensemblgenomes.ebi.ac.uk/pub/plants/release-59/fasta/oryza_punctata/pep/Oryza_punctata.Oryza_punctata_v1.2.pep.all.fa.gz

### Evaluation and benchmark

The CAFA-evaluator (https://github.com/BioComputingUP/CAFA-evaluator) was used to calculate the weighted precision (the proportion of correct positive predictions out of all positive predictions), recall (the proportion of positive predictions made out of all positive examples in the dataset), and F-score for each GO domain independently. OmicsBox/BLAST2GO was used as a benchmark. To avoid sample biases, a sample of 1,000 genes was extracted in triplicate from *A. thaliana*. Welch’s t-test was conducted for each GO main domain to compare the performance of the tools. The GO annotation of *A. thaliana* from the Gene Ontology Consortium website was considered the gold standard annotation for the purpose of our comparison.

## Results and discussion

### Ortholog enrichment

Among plant species, 50.9% of UniProt SwissProt accessions (22,658 accessions) and 14.2% of UniProt Trembl accessions (2,706,481 accessions) have been annotated with OrthoDB OG. After enrichment of UniProt-Plants with OrthoDB OG ID (for simplicity, ODB-ID), using USEARCH 64-bit, the ODB-IDs increased from 14.3% to 48.8% (Figure 1A). The enriched ODB-IDs, using the sequence alignment approach against OrthoDB, were compared with the ODB-IDs originally annotated in UniProtKB. The results showed that exactly the same ODB-IDs were present in 99.6% of the cases among the 2,729,139 accessions (14.3%) originally annotated with ODB-IDs. This finding instilled confidence in the enrichment approach as a feasible method for inferring orthologs on a large scale.

### Annotation per protein existence level

To demonstrate Hayai-Annotation v3’s capabilities regarding protein existence levels, we annotated and compared three species: *A. thaliana, O. sativa*, and *O. punctata* (Table 1). As expected, model organisms or species that are more extensively studied exhibited greater numbers of annotated genes at a protein level (PE 1).

**Table 1.** Comparison of the number of annotated genes based on the protein existence level as defined by UniProt.

|  | Protein Existence (PE) Level |  |  |  |  |
| --- | --- | --- | --- | --- | --- |
|  | 1<br>(Protein) | 2<br>(Transcript) | 3<br>(Homology) | 4<br>(Predicted) | 5<br>(Uncertain) |
| *A. thaliana* | 12,562 | 6,765 | 2,619 | 5,499 | 67 |
| *O. sativa* | 7,767 | 6,133 | 7,947 | 20,764 | 7 |
| *O. punctata* | 81 | 90 | 14,893 | 25,922 | 0 |

### GO enrichment evaluation

Using the CAFA-evaluator, weighted precision, recall, and F scores were calculated for each GO domain independently. The results showed no statistically significant difference at the 0.05 significance level in the MF and BP domains (Table 2, Supplementary File Table S2). Hayai-Annotation v3 exhibited superior performance in the CC domain (P = 0.002305).

**Table 2.**
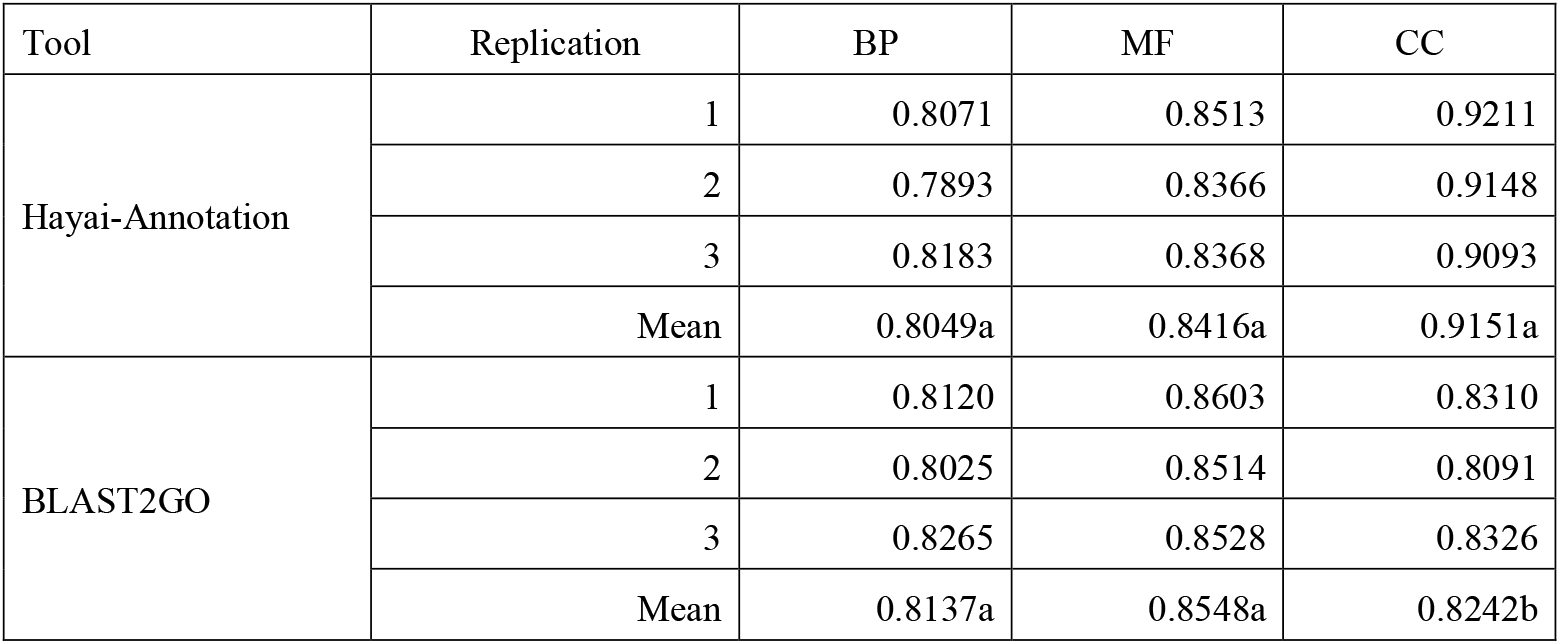
Weighted F calculated using the CAFA-evaluator for a sample of 1,000 genes from *A. thaliana* with three replications. Letters (a and b) represent the statistical differences determined by Welch’s t-test (alpha = 0.05).

In our previous publications about Hayai-Annotation Plants, we considered only child GO terms and disregarded parental terms to evaluate the performance of all tested software tools. In response to observations by Van Bel and Vandepoele (2020)^16^, we realized that the comparison of tools in our previous publications was insufficient. Therefore, we utilized the AUC-PR comparison data described in our earlier paper as a response to their comments and submitted it to the journal that had published the report by Van Bel and Vandepoele (2020). Unfortunately, we were unable to secure the opportunity to publish the results of this analysis in that journal. Consequently, despite our reservations about the situation, our paper that Van Bel and Vandepoele (2020) commented on was retracted. We therefore published the results of benchmarking the original version of Hayai-Annotation Plants using AUC-PR and the CAFA-evaluator, proving that Hayai-Annotation possesses sufficient accuracy as a gene function annotation tool^15^. The results of that study indicate that the functional annotation accuracy of Hayai-Annotation v3 is also sufficiently high.

### Co-occurrence and network analysis

We propose a method to gain a different perspective on genome evolution and adaptation by examining the co-occurrence of ODB IDs with GO MF and of ODB IDs with GO BP. Gene duplications, functional redundancy, or specialization may ensue, which is especially crucial in plants due to their immobility. Therefore, although a set of genes may share the same ODB ID, they could be involved in distinct MFs or BPs. Furthermore, convergent evolution, where the same MF may appear in different ODB IDs, is a significant evolutionary occurrence. To grasp the broader context, we propose the analysis of co-occurrence IDs between ODB and GO (MF and BP).

The co-occurrence tables and network analysis seemed particularly interesting when we compared different species within the same genus. To demonstrate some of the potential of this analysis, we analyzed the *Oryza sativa* Japonica Group, cultivar Nipponbare, with the wild-type *O. punctata*. The resulting networks, developed using the top 100 co-occurrences and considering the gene count for each ODB ID–GO (MF and BP), were compared.

In the wild type, the GO MF “ATP binding” (GO:0005524) showed more diverse ODB IDs compared to domesticated rice, while the MF “RNA binding” (GO:0003723) and “ADP binding” (GO:0043531) were present only in the wild type (Suppl. File Figure S1). In the cultivated rice, the MF “nutrient reservoir activity” (GO:0045735), besides the ODB “Legumin A” (925447at33090), showed 25 genes with the ODB “Bifunctional inhibitor/plant lipid transfer protein/seed storage helical domain” (147891at33090, referred to as seed storage in Suppl. File Figure S2). Also, in the MF “ATP binding” the cultivated rice showed the ODB “LEAF RUST 10 DISEASE-RESISTANCE LOCUS RECEPTOR-LIKE PROTEIN KINASE-like 2.1” (177763at33090). Leaf Rust Kinase 10-like proteins are known to participate in plant growth and stress responses, particularly in controlling the time of flowering and defense responses against pathogens^17^. Different patterns of ODB IDs were present under the “DNA binding” (GO:0003677) and “DNA-binding transcription factor activity” (GO:0003700) MFs. The co-occurrence of ODB with the latter GO terms revealed that in the wild type, 9 genes were associated with “Floral homeotic protein APETALA 2” (103234at33090), while in the cultivated rice, 5 genes were linked to it. The ODB “AP2-like ethylene-responsive transcription factor AIL5” (695999at33090) was found in 14 genes in *O. sativa* and in 1 gene in the wild type (101433at33090). The complete data can be found in the co-occurrence tables (Supplementary Table S1).

The network analysis of ODB–GO BP showed a different pattern for the BP “phosphorylation” (GO:0016310). In the wild type, more genes were linked to “protein kinase C-activating G protein-coupled receptor signaling pathway” (GO:0007205) and “Signal transduction” (GO:0007165), while in cultivated rice, the related BPs were “Intracellular signal transduction” (GO:0035556) and “Cell surface receptor signaling pathway” (GO:0007166) (Suppl. File Figure S3). In the wild type, the top ODB was “serine/threonine-protein kinase OXI1-like” (988924at33090), whereas in the cultivated rice, it was “LEAF RUST 10 DISEASE-RESISTANCE LOCUS RECEPTOR-LIKE PROTEIN KINASE-like 2.1” (177763at33090) (Suppl. File Figure S4). Also, the wild type showed that the BP “microtubule-based movement” (GO:0007018) co-occurred with three ODB IDs related to “Kinesin-like protein” (114566at33090, 175388at33090, and 23898at33090).

Microtubules play a critical role in defining plant cell shape by directing cell expansion, a crucial process for adapting to changing environmental conditions^18^. The mutation of kinesin genes resulted in disorganized cortical microtubules and abnormal cell shapes^19^. This phenotypic plasticity is essential for environmental adaptation, and it is a trait pursued in plant breeding programs to enable cultivation in diverse environmental conditions. However, in terms of crop management for commercial purposes, phenotypic variability is considered the least desirable characteristic.

Therefore, we showed that Hayai-Annotation v3 was able to accurately predict GO terms and to assign orthologs by using two methods, giving priority to OrthoLoger, the official tool developed by OrthoDB. We also demonstrated that the network analysis, based on the co-occurrence tables of ODB IDs with GO (MF and BP), provided as simple way to visualize the traits selected by a species, whether through natural selection to maintain diversity or in breeding programs.

## Conclusion

Bioinformatics techniques for genome sequence analysis have advanced alongside the advancement of computer technology. Historically, emphasis has been placed on genome assembly techniques, but combinations like PacBio’s HiFi reads and assembly tools such as Hifiasm^20^ now allow for the generation of near-chromosome-level scaffolds at relatively low cost. In contrast, gene prediction technologies had seen little advancement for a long time. However, the emergence of tools such as Helixer^21^, which leverages deep learning on GPUs for gene prediction, has significantly reduced the time and cost associated with gene prediction. There is a demand for tools that can perform rapid calculations and provide cross-species functional annotation. For example, Plant GARDEN, a plant genome portal site, re-annotates all gene information in its database using Hayai-Annotation to compare gene sequence information generated by various methods^22^. Hayai-Annotation v3 not only retains the convenience of previous versions but also enhances the functionality of ortholog analysis, allowing for evolutionary insights from gene sequences. Hayai-Annotation v3 is expected to contribute significantly to the future development of plant genome analysis.

## Acknowledgements

This work was supported by research funds from the National Institute of Genetics and the Kazusa DNA Research Institute.

## Data Availability

The functionality of Hayai-Annotation Plants is publicly available through the following site (https://github.com/aghelfi/HayaiAnnotation).

## Notes

https://github.com/aghelfi/HayaiAnnotation

### Competing Interest Statement

The authors have declared no competing interest.

## References

1. Marks, R. A., Hotaling, S., Frandsen, P. B., and VanBuren, R. 2021, Representation and participation across 20 years of plant genome sequencing. Nature Plants, 7, 1571–8.

2. Conesa, A., and Götz, S. 2008, Blast2GO: A comprehensive suite for functional analysis in plant genomics. Int. J. Plant Genomics, 2008, 619832.

3. Schwacke, R., Ponce-Soto, G. Y., Krause, K., et al. 2019, MapMan4: A Refined Protein Classification and Annotation Framework Applicable to Multi-Omics Data Analysis. Mol. Plant, 12, 879–92.

4. Chen, T.-W., Gan, R.-C., Fang, Y.-K., et al. 2017, FunctionAnnotator, a versatile and efficient web tool for non-model organism annotation. Sci. Rep., 7, 1–9.

5. Altschul, S. F., Gish, W., Miller, W., Myers, E. W., and Lipman, D. J. 1990, Basic local alignment search tool. J. Mol. Biol., 215, 403–10.

6. Buchfink, B., Reuter, K., and Drost, H.-G. 2021, Sensitive protein alignments at tree-of-life scale using DIAMOND. Nat. Methods, 18, 366–8.

7. Zdobnov, E. M., and Apweiler, R. 2001, InterProScan--an integration platform for the signature-recognition methods in InterPro. Bioinformatics, 17, 847–8.

8. Eddy, S. R. 2011, Accelerated Profile HMM Searches. PLoS Comput. Biol., 7, e1002195.

9. Frith, M. C., Hamada, M., and Horton, P. 2010, Parameters for accurate genome alignment. BMC Bioinformatics, 11, 80.

10. Altschul, S. F., Madden, T. L., Schäffer, A. A., et al. 1997, Gapped BLAST and PSI-BLAST: a new generation of protein database search programs. Nucleic Acids Res., 25, 3389–402.

11. Kuznetsov, D., Tegenfeldt, F., Manni, M., et al. 2022, OrthoDB v11: annotation of orthologs in the widest sampling of organismal diversity. Nucleic Acids Res., 51, D445–51.

12. Emms, D. M., and Kelly, S. 2019, OrthoFinder: phylogenetic orthology inference for comparative genomics. Genome Biol., 20, 238.

13. Li, L., Stoeckert, C. J., Jr, and Roos, D. S. 2003, OrthoMCL: identification of ortholog groups for eukaryotic genomes. Genome Res., 13, 2178–89.

14. Ghelfi, A., Shirasawa, K., Hirakawa, H., and Isobe, S. 2018, November 20, Hayai-Annotation Plants: an ultra-fast and comprehensive gene annotation system in plants. bioRxiv, p. 473488.

15. Ghelfi, A., Shirasawa, K., and Isobe, S. 2023, September 12, Benchmarking Hayai-Annotation Plants: A Re-evaluation Using Standard Evaluation Metrics. bioRxiv, p. 2023.09.08.556781.

16. Van Bel, M., and Vandepoele, K. 2020, Comment on ‘Hayai-Annotation Plants: an ultrafast and comprehensive functional gene annotation system in plants’: the importance of taking the GO graph structure into account. Bioinformatics, 36, 5558–60.

17. Gandhi, A., and Oelmüller, R. 2023, Emerging Roles of Receptor-like Protein Kinases in Plant Response to Abiotic Stresses. Int. J. Mol. Sci., 24.

18. Yang, B., Wendrich, J. R., De Rybel, B., Weijers, D., and Xue, H.-W. 2020, Rice microtubule-associated protein IQ67-DOMAIN14 regulates grain shape by modulating microtubule cytoskeleton dynamics. Plant Biotechnol. J., 18, 1141–52.

19. Zhang, M., Zhang, B., Qian, Q., et al. 2010, Brittle Culm 12, a dual-targeting kinesin-4 protein, controls cell-cycle progression and wall properties in rice. Plant J., 63, 312–28.

20. Cheng, H., Concepcion, G. T., Feng, X., Zhang, H., and Li, H. 2021, Haplotype-resolved de novo assembly using phased assembly graphs with hifiasm. Nat. Methods, 18, 170–5.

21. Stiehler, F., Steinborn, M., Scholz, S., Dey, D., Weber, A. P. M., and Denton, A. K. 2020, Helixer: cross-species gene annotation of large eukaryotic genomes using deep learning. Bioinformatics, 36, 5291–8.

22. Ichihara, H., Yamada, M., Kohara, M., et al. 2023, Plant GARDEN: a portal website for cross-searching between different types of genomic and genetic resources in a wide variety of plant species. BMC Plant Biol., 23, 391.

