## Supplementary_File_S1 for "Hayai-Annotation v3.0: A functional gene prediction tool that integrates orthologs and gene ontology for network analysis"

Corresponding Author

* 'To whom correspondence should be addressed'

Table S1 Complete co-occurrence tables of OrthoDB IDs and GO (MF and BP) for wild rice (tabs 1 and 2) and *O. sativa* (tabs 3 and 4). Filename: Table_S1.xlsx

Table S2 Benchmark of GO enrichments using CAFA-evaluation, weighted PR, RC, and F, with *A. thaliana*, comparing Hayai-Annotation v3 and BLAST2GO. Replications 1-3 (A, B, C).

(A)

| filename | ns | tau | n | pr_w | rc_w | f_w |
|---|---|---|---|---|---|---|
| blast2go_rep1.tsv | biological_process | 0.001 | 550 | 0.9574 | 0.7050 | 0.8120 |
| blast2go_rep1.tsv | cellular_component | 0.001 | 612 | 0.9848 | 0.7188 | 0.8310 |
| blast2go_rep1.tsv | molecular_function | 0.001 | 605 | 0.9813 | 0.7659 | 0.8603 |
| hayai_rep1.tsv | biological_process | 0.001 | 538 | 0.9627 | 0.6948 | 0.8071 |
| hayai_rep1.tsv | cellular_component | 0.001 | 648 | 0.9851 | 0.8649 | 0.9211 |
| hayai_rep1.tsv | molecular_function | 0.001 | 579 | 0.9928 | 0.7451 | 0.8513 |

(B)

| filename | ns | tau | n | pr_w | rc_w | f_w |
|---|---|---|---|---|---|---|
| blast2go_rep2.tsv | biological_process | 0.001 | 531 | 0.963 | 0.6879 | 0.8025 |
| blast2go_rep2.tsv | cellular_component | 0.001 | 570 | 0.9854 | 0.6863 | 0.8091 |
| blast2go_rep2.tsv | molecular_function | 0.001 | 594 | 0.9782 | 0.7538 | 0.8514 |
| hayai_rep2.tsv | biological_process | 0.001 | 512 | 0.9713 | 0.6647 | 0.7893 |
| hayai_rep2.tsv | cellular_component | 0.001 | 620 | 0.9894 | 0.8507 | 0.9148 |
| hayai_rep2.tsv | molecular_function | 0.001 | 567 | 0.9909 | 0.7238 | 0.8366 |

(C)

| filename | ns | tau | n | pr_w | rc_w | f_w |
|---|---|---|---|---|---|---|
| blast2go_rep3.tsv | biological_process | 0.001 | 545 | 0.9704 | 0.7198 | 0.8265 |
| blast2go_rep3.tsv | cellular_component | 0.001 | 588 | 0.9821 | 0.7226 | 0.8326 |
| blast2go_rep3.tsv | molecular_function | 0.001 | 593 | 0.9848 | 0.7521 | 0.8528 |
| hayai_rep3.tsv | biological_process | 0.001 | 528 | 0.9777 | 0.7036 | 0.8183 |
| hayai_rep3.tsv | cellular_component | 0.001 | 615 | 0.989 | 0.8415 | 0.9093 |
| hayai_rep3.tsv | molecular_function | 0.001 | 575 | 0.9914 | 0.7239 | 0.8368 |

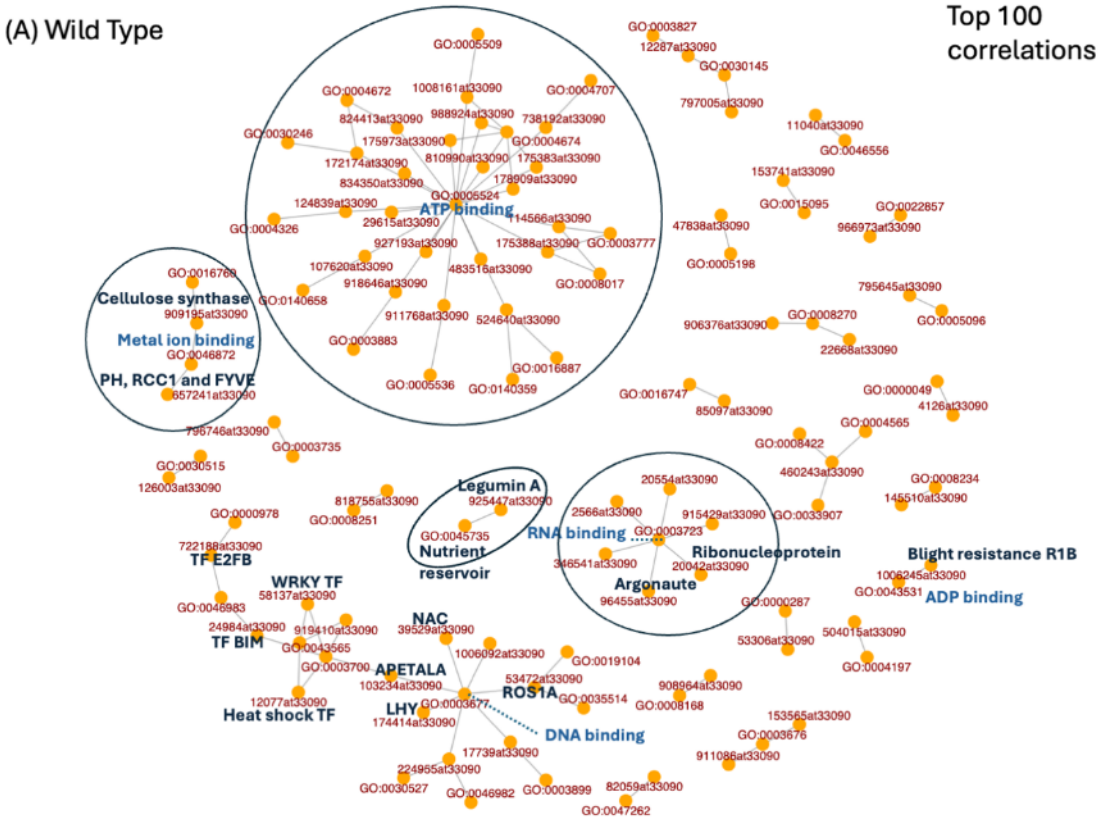
(A) Wild Type
Top 100 correlations
ATP binding
Cellulose synthase
Metal ion binding
PH, RCC1 and FYVE
Legumin A
Nutrient reservoir
RNA binding
Ribonucleoprotein
Argonaute
Blight resistance R1B
ADP binding
TF E2FB
WRKY TF
TF BIM
NAC
APETALA
ROS1A
LHY
Heat shock TF
DNA binding
GO:0005509
1008161at33090
GO:0004672
GO:0004707
824413at33090
988924at33090
738192at33090
GO:0030246
175973at33090
GO:0004674
172174at33090
810990at33090
175383at33090
834350at33090
178909at33090
GO:0005524
124839at33090
114566at33090
GO:0004326
29615at33090
927193at33090
175388at33090
GO:0003777
107620at33090
483516at33090
GO:0008017
918646at33090
GO:0140658
911768at33090
524640at33090
GO:0003883
GO:0016887
GO:0005536
GO:0140359
GO:0016760
909195at33090
GO:0046872
657241at33090
GO:0003827
12287at33090
GO:0030145
797005at33090
11040at33090
GO:0046556
153741at33090
GO:0015095
GO:0022857
47838at33090
966973at33090
GO:0005198
795645at33090
GO:0005096
906376at33090
GO:0008270
22668at33090
GO:0016747
85097at33090
GO:0000049
4126at33090
GO:0004565
GO:0008422
460243at33090
GO:0008234
145510at33090
GO:0033907
796746at33090
GO:0003735
GO:0030515
126003at33090
818755at33090
GO:0008251
925447at33090
GO:0045735
20554at33090
2566at33090
915429at33090
GO:0003723
346541at33090
20042at33090
96455at33090
GO:0000287
1006245at33090
GO:0043531
GO:0000978
722188at33090
58137at33090
GO:0046983
919410at33090
24984at33090
GO:0043565
GO:0003700
39529at33090
1006092at33090
GO:0019104
53472at33090
103234at33090
GO:0003677
GO:0035514
908964at33090
GO:0008168
12077at33090
174414at33090
53306at33090
504015at33090
GO:0004197
153565at33090
GO:0003676
911086at33090
17739at33090
224955at33090
GO:0030527
GO:0046982
GO:0003899
82059at33090
GO:0047262

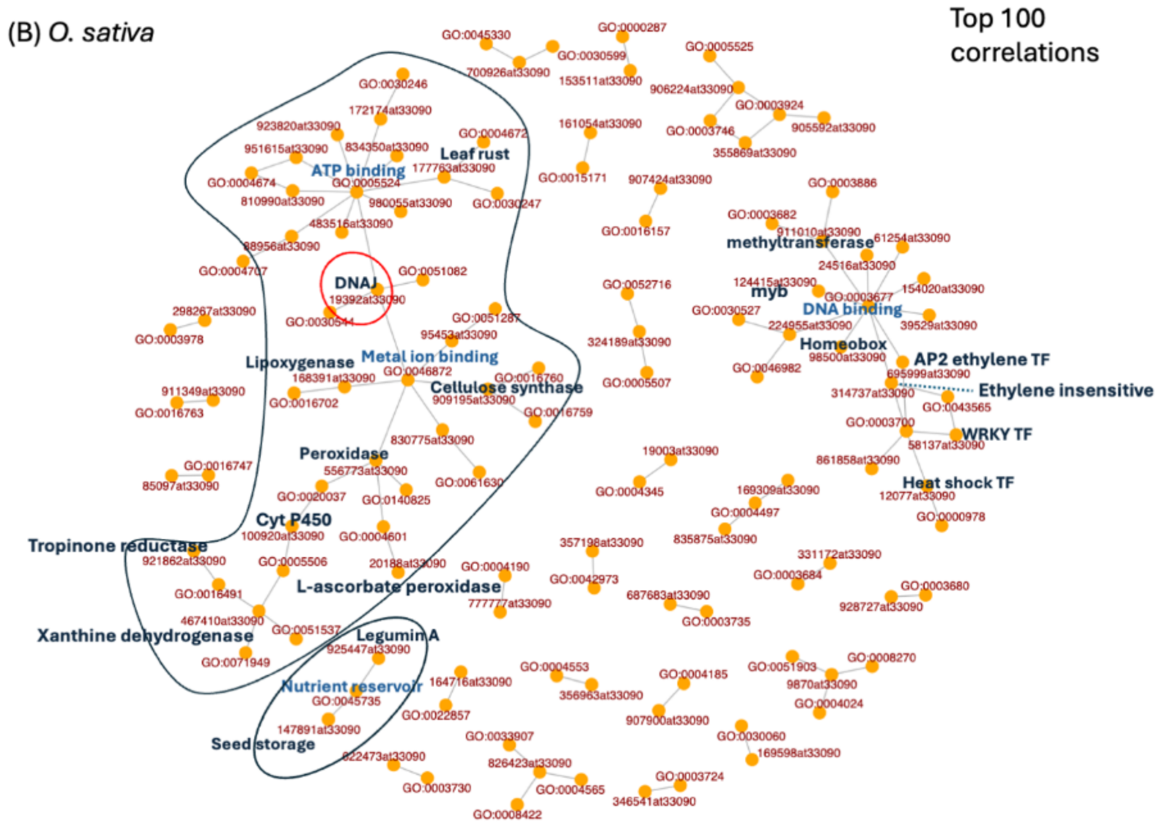


Figure S1. Network analysis performed on the top 100 co-occurrences of ODB IDs and GO molecular function. (A) wild type; (B) cultivated rice.

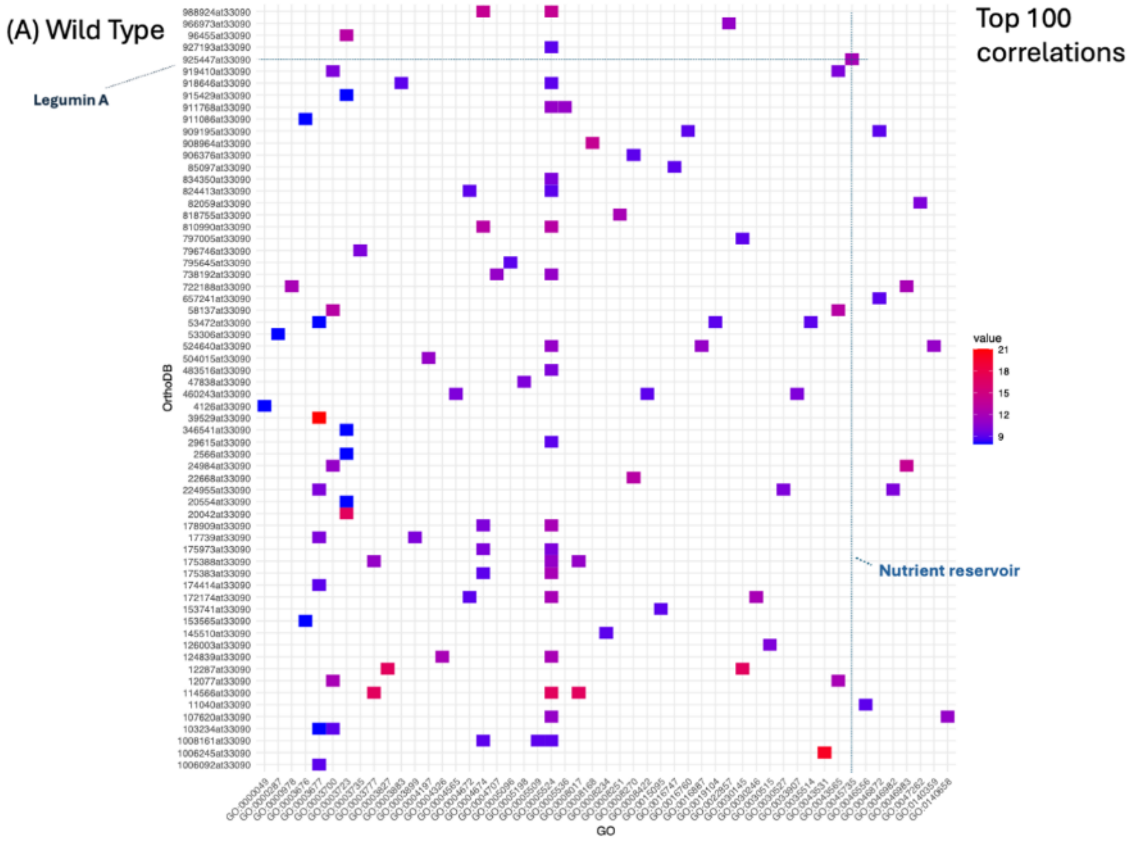

(A) Wild Type
Top 100 correlations
Legumin A
Nutrient reservoir
OrthoDB
GO
value
21
18
15
12
9
988924at33090
966973at33090
96455at33090
927193at33090
925447at33090
919410at33090
918646at33090
915429at33090
911768at33090
911086at33090
909195at33090
908964at33090
906376at33090
85097at33090
834350at33090
824413at33090
82059at33090
818755at33090
810990at33090
797005at33090
796746at33090
795645at33090
738192at33090
722188at33090
657241at33090
58137at33090
53472at33090
53306at33090
524640at33090
504015at33090
483516at33090
47838at33090
460243at33090
4126at33090
39529at33090
346541at33090
29615at33090
2566at33090
24984at33090
22668at33090
224955at33090
20554at33090
20042at33090
178909at33090
17739at33090
175973at33090
175388at33090
175383at33090
174414at33090
172174at33090
153741at33090
153565at33090
145510at33090
126003at33090
124839at33090
12287at33090
12077at33090
114566at33090
11040at33090
107620at33090
103234at33090
1008161at33090
1006245at33090
1006092at33090

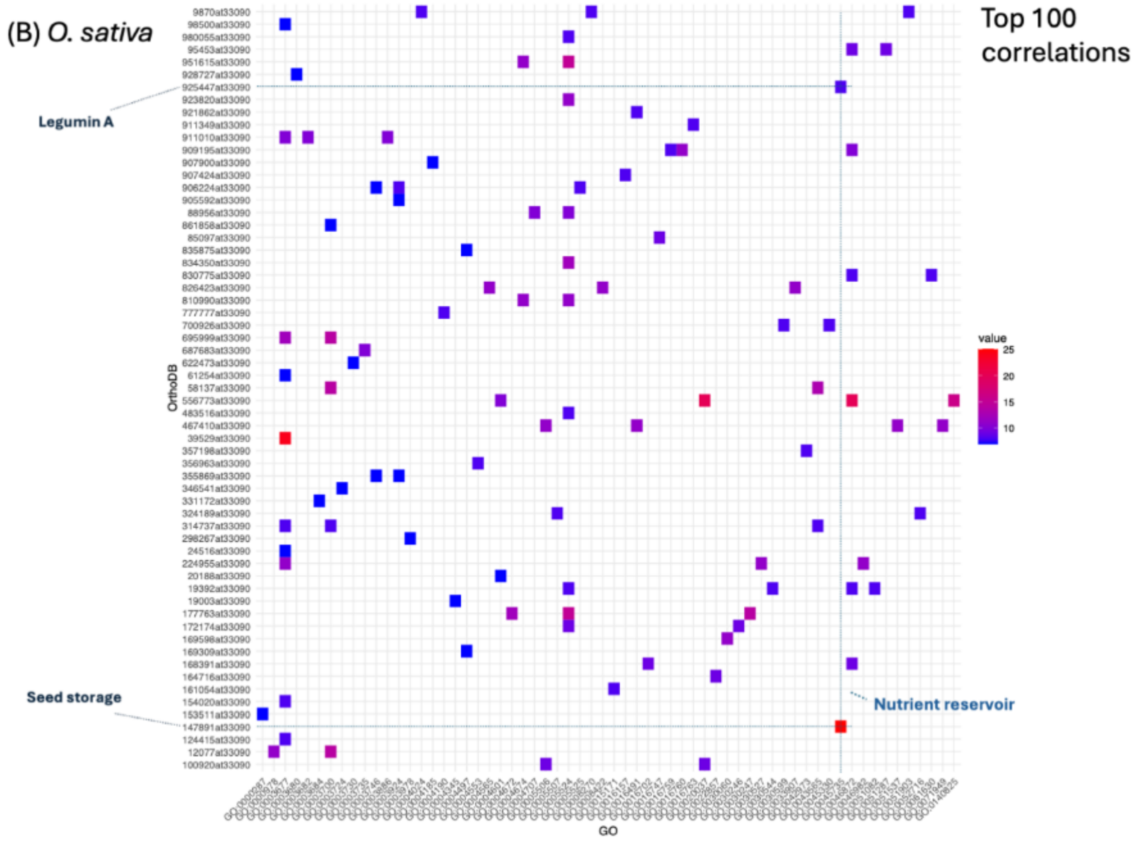


Figure S2. Heatmap analysis performed on the top 100 co-occurrences of ODB IDs and GO molecular function. (A) wild type; (B) cultivated rice.

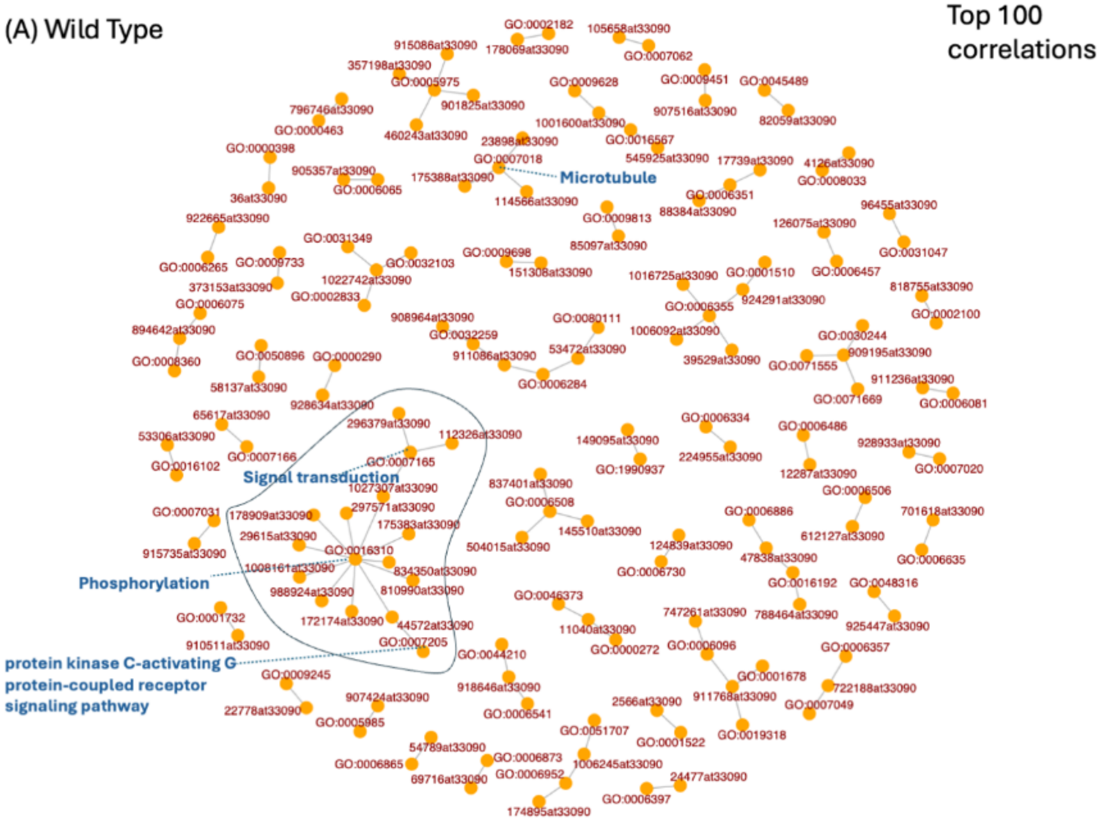
(A) Wild Type
Top 100 correlations
Microtubule
Signal transduction
Phosphorylation
protein kinase C-activating G protein-coupled receptor signaling pathway

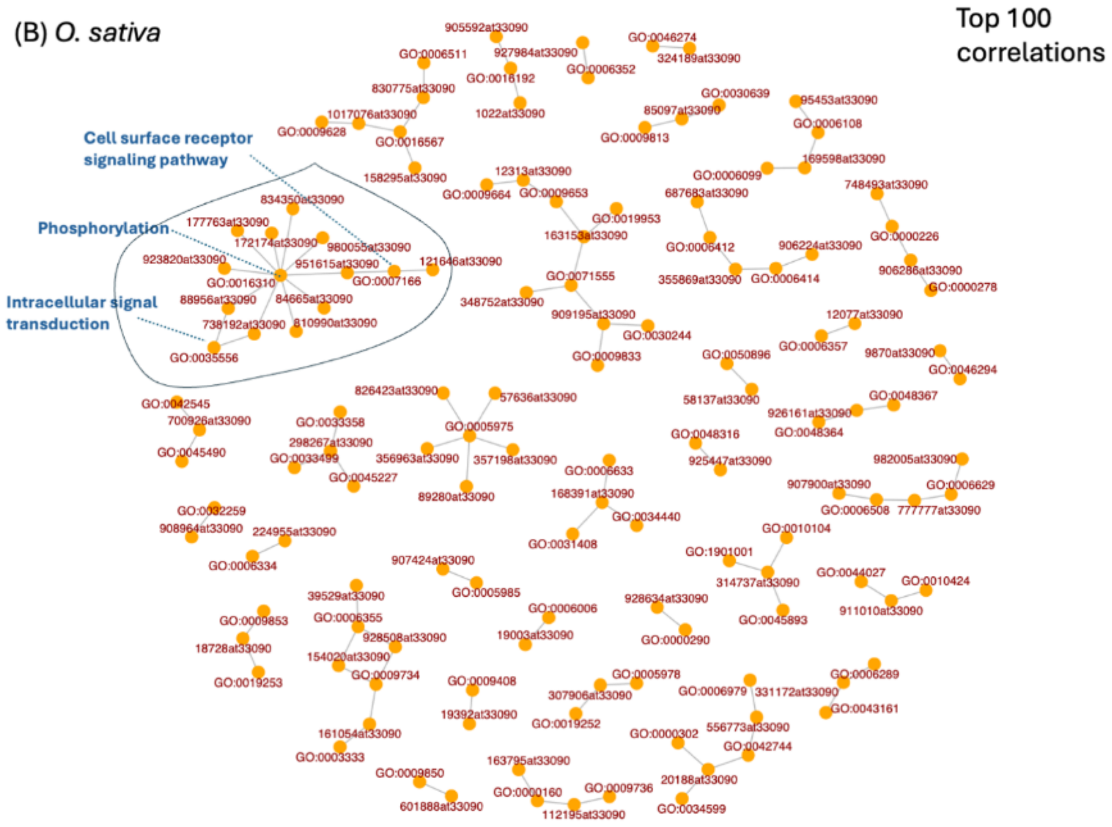


Figure S3. Network analysis performed on the top 100 co-occurrences of ODB IDs and GO biological process. (A) wild type; (B) cultivated rice.

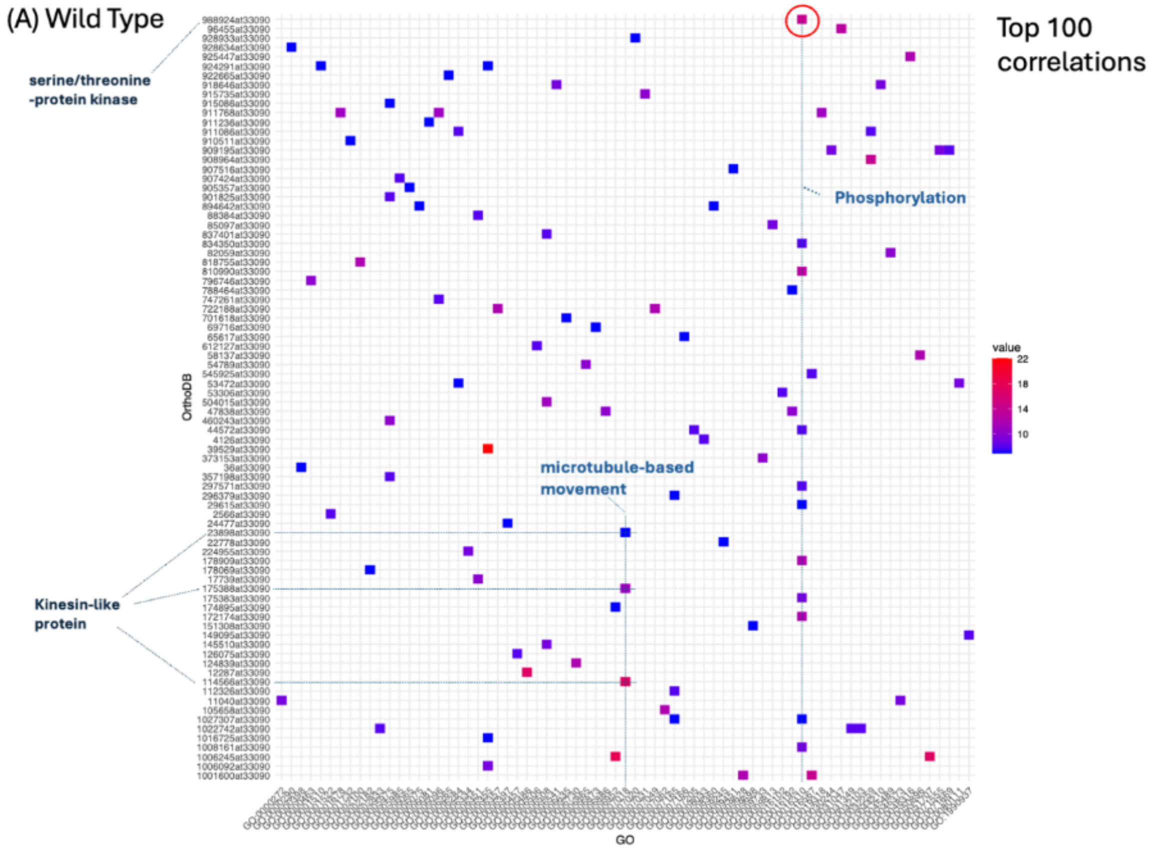
(A) Wild Type
Top 100 correlations
serine/threonine -protein kinase
Phosphorylation
microtubule-based movement
Kinesin-like protein
value
22
18
14
10
OrthoDB
GO

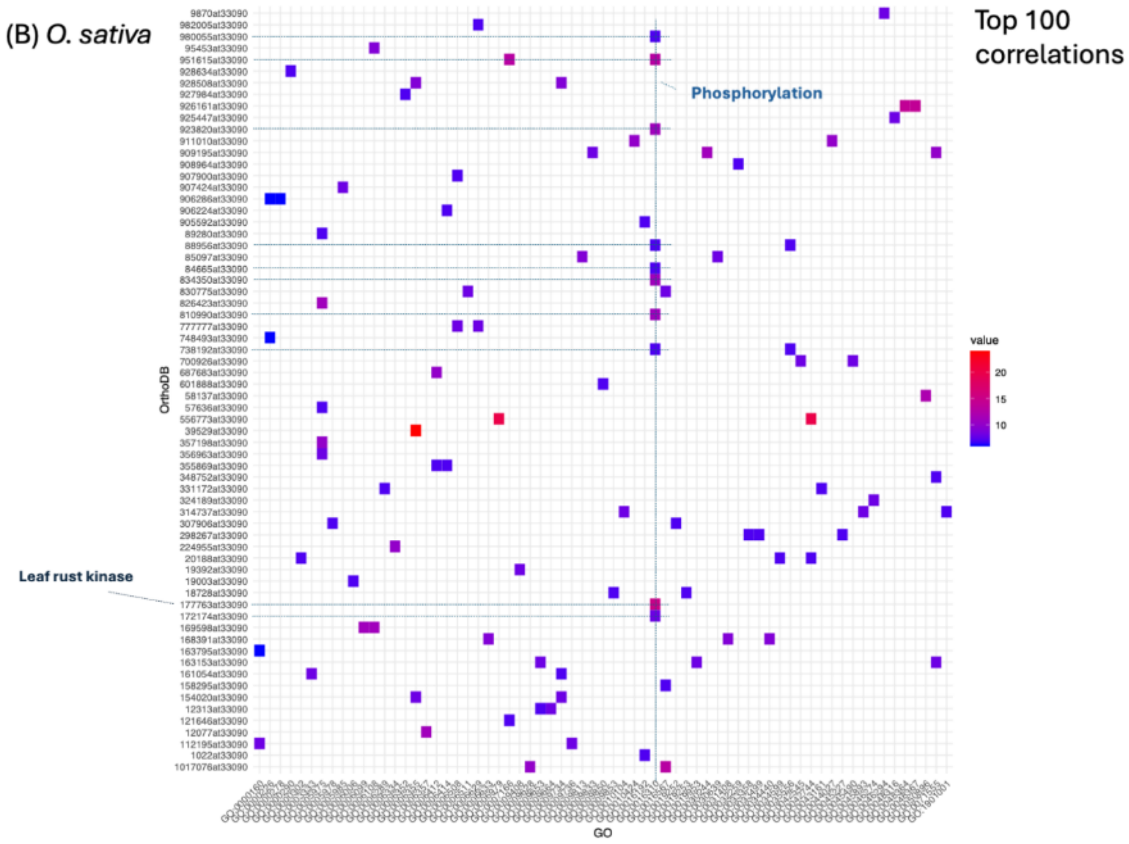


Figure S4. Heat analysis performed on the top 100 co-occurrences of ODB IDs and GO biological process. (A) wild type; (B) cultivated rice.

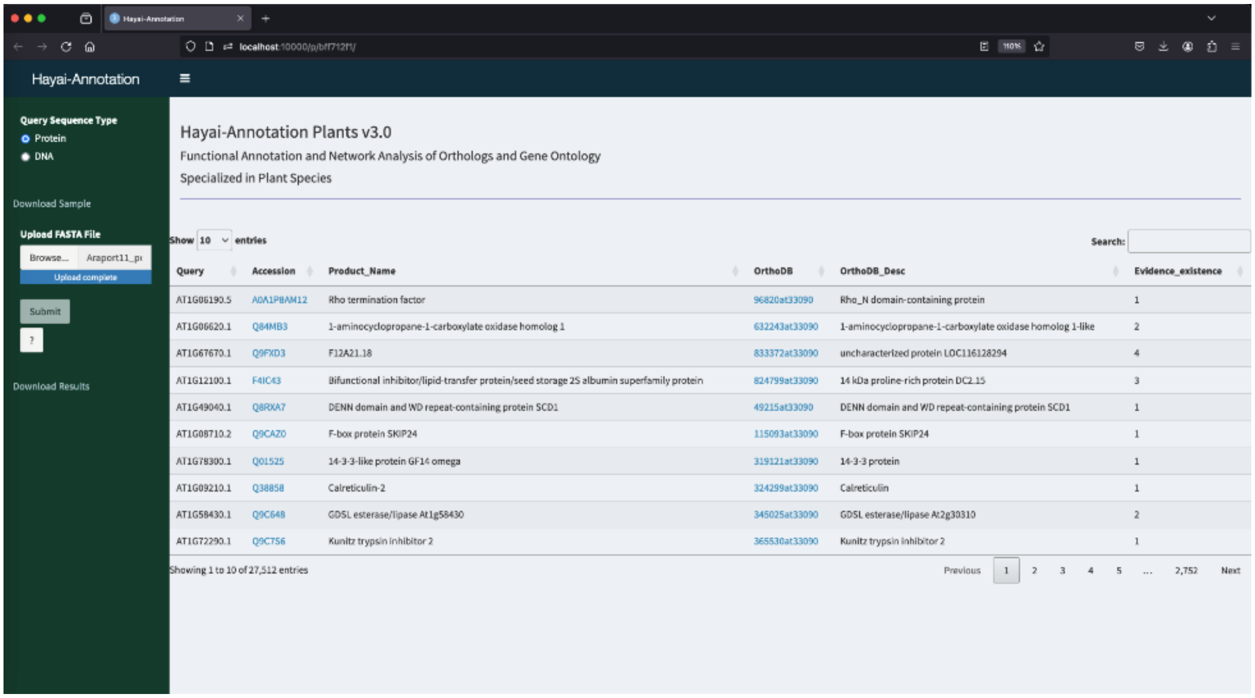


Figure S5. Hayai-Annotation v3 interface.

Hayai-Annotation v3 generates an output file named 'output_HayaiAnnotation_v2.zip'. It comprises 7 tables (TSV format) and 8 graphics (PDF format). The files provided correspond to the full annotation of *A. thaliana*.

Hayai_annotation_v3.0.tsv is the main table and contains the full annotations from both methods, DIAMOND and Orthomapper (OrthoLoger).

Four tables aggregate the results for each GO domain (MF, BP, CC) and InterPro annotation layers; they are named, respectively, 'Hayai_annotation_GO_MF.tsv', 'Hayai_annotation_GO_BP.tsv', 'Hayai_annotation_GO_CC.tsv' and 'Hayai_annotation_Interpro.tsv'.

Two tables show the co-occurrences of ODB ID and GO (MF and BP, independently), named 'Correlations_OrthoDB_GO_MF.tsv' and 'Correlations_OrthoDB_GO_BP.tsv'. The count corresponds to the number of genes for each co-occurrence.

Four graphics are generated, two regarding the network and two the heatmap, based on the top 100 co-occurrences of ODB ID and GO (MF and BP, independently). The filenames are: 'Graph_Network_OrthoDB_GO_MF.pdf', 'Graph_Network_OrthoDB_GO_BP.pdf', 'Graph_Heatmap_OrthoDB_GO_MF.pdf' and 'Graph_Heatmap_OrthoDB_GO_BP.pdf'.

Using the top 50 counts of the aggregated results, four graphics are generated for the distribution of GO (MF, BP and CC) and InterPro.
